# Robust, versatile DNA FISH probes for chromosome-specific repeats in *Caenorhabditis elegans* and *Pristionchus pacificus*

**DOI:** 10.1101/2022.01.25.477430

**Authors:** Renzo S. Adilardi, Abby F. Dernburg

**Affiliations:** Department of Molecular and Cell Biology, University of California, Berkeley, Berkeley, CA USA; Howard Hughes Medical Institute, Chevy Chase, MD USA; Biological Systems and Engineering Division, Lawrence Berkeley National Laboratory, Berkeley, CA USA; California Institute for Quantitative Biosciences, Berkeley, CA USA

## Abstract

Repetitive DNA sequences are useful targets for chromosomal fluorescence in situ hybridization (FISH). We analyzed recent genome assemblies of *Caenorhabditis elegans* and *Pristionchus pacificus* to identify tandem repeats with a unique genomic localization. Based on these findings, we designed and validated sets of oligonucleotide probes for each species targeting at least one locus per chromosome. These probes yielded reliable fluorescent signals in different tissues and can easily be combined with immunolocalization of cellular proteins. Synthesis and labeling of these probes are highly cost-effective and require no hands-on labor. The methods presented here can be easily applied in other model and non-model organisms with a sequenced genome.

## INTRODUCTION

Fluorescence in situ hybridization (FISH) targeting chromosomal DNA has long been a powerful cytological tool to study chromosomes and nuclear architecture in fixed samples (Pardue and Gall 1969; Bauman *et al*. 1980). Successful application of this technique depends on design and synthesis of suitable labeled probes, combined with development of fixation and hybridization methods appropriate to address specific biological questions.

While many experiments require custom-designed probes to specific genomic regions of interest, a wide range of questions can also be addressed using probes to tandemly repeated sequences. Such repeats are often a major component of eukaryotic genomes, and typically concentrate at telomeric and pericentromeric regions. Multigene families such as ribosomal RNA genes (rDNA) and tandem repeats of varying lengths (satellite DNA) are commonly used as FISH targets in studies of genome organization, karyotype evolution, or chromosome segregation (e.g., Dernburg *et al*. 1996; Nguyen *et al*. 2010; Heckmann *et al*. 2013; Ruiz-Ruano *et al*. 2016). These and other repetitive sequences that are restricted to one or a few genomic loci are particularly valuable for cytological studies.

Several technical features make tandemly-repeated sequences favorable targets for DNA FISH experiments. Probes that detect such sequences are easy to generate, since one or a few short, synthetic oligonucleotides can hybridize to many clustered targets. Short DNA fragments are ideal FISH probes, particularly for whole-mount samples, as they readily diffuse through tissue fixed with moderate concentrations of formaldehyde (i.e., 2-5%), which is sufficient to preserve morphology during the denaturation steps required for DNA FISH, usually accomplished with a combination of heat and formamide. DNA oligonucleotides can readily be labeled with fluorophores or haptens either during chemical synthesis or afterwards, using enzymatic methods. Synthesis of probes with modified backbones, such as Locked Nucleic Acid (LNA) or Peptide Nucleic Acid (PNA) chemistries, can further enhance detection by stabilizing probe-target hybrid molecules. Detection of high copy-number sequences is inherently robust due to favorable hybridization kinetics arising from the high effective concentration of the target. Thus, repetitive probes typically produce more intense and reliable signals than probes targeting a single-copy sequence spanning a similar genomic interval. This is particularly advantageous in well-preserved whole-mount cells or tissues, where hybridization of the probe to the target competes with reannealing of denatured chromosomal DNA during the hybridization process. While synthetic oligonucleotide probes can also be designed to detect single-copy sequences (e.g., Oligopaints; Beliveau *et al*. 2012), they must be used at a much higher total concentration, at least proportional to the sequence complexity. This requires significant effort and cost to amplify complex probes from synthetic libraries, which typically yield only low femtomolar quantities of each oligo (Schmidt *et al*. 2015)

The genomes of *Caenorhabditis elegans* and most other sequenced nematodes are compact and have low repeat content compared to many other eukaryotes (Rödelsperger *et al*. 2013). Consequently, only a few repetitive FISH probes have been described for *C. elegans*: these include probes against the telomere repeats (TTAGGC)n, the large ribosomal RNA gene cluster (8S, 5.8S, and 26S rDNA) on chromosome I, the tandemly repeated 5S rDNA locus on chromosome V, and two clustered repeats on the X chromosome (Albertson 1984; Dernburg *et al*. 1998; Phillips *et al*. 2005; Ferreira *et al*. 2013).

A new reference *C. elegans* genome was recently assembled based on long-read sequence data for the VC2010 strain (derived from N2), which increased the accuracy of repeat annotations (Yoshimura *et al*. 2019). This study reported an additional 1.8Mb of tandem repeats and other duplications relative to previous assemblies, a significant difference in a genome comprising ~100 Mb. Similar telomere-to-telomere genome assemblies with accurately annotated repetitive sequences are becoming the norm for many non-traditional model species, as well as for specific isolates or strains of highly studied organisms.

Leveraging current genome assemblies, we searched the genomes of *C. elegans* and its distant relative, *Pristionchus pacificus*, for tandem repeats suitable for DNA FISH in whole-mount tissues. We selected candidate repeats restricted to a single major genomic locus, designed simple, inexpensive FISH probes, and tested their performance in different tissues. Through this straightforward approach we have designed and validated a set of probes that include at least one locus per chromosome for each species. The criteria we used proved to be highly reliable for target selection and probe design and can be easily extended to design FISH probes for other model and non-model organisms.

## MATERIALS AND METHODS

### Tandem repeat analysis

Chromosome-level genome assemblies from *C. elegans* (“VC2010”, Yoshimura *et al*. 2019), and *P. pacificus* (“El Paco”, Rödelsperger *et al*. 2017) were analyzed using Tandem Repeat Finder v4.09 (Benson 1999). A maximum periodicity of 200 bp and default parameters were used (match = 2, mismatch = 7, indels = 7, minimum alignment score to report repeat = 50). The total span of each repeat was calculated using the genomic indices. The output data for each chromosome were filtered to select repeats spanning more than 5 kb. These candidate repeats were analyzed using BLAST against the reference genomes to characterize their distributions. For longer tandem repeats (monomer >40bp), the repetitive motif was split into nonoverlapping subsequences, each of which was reanalyzed by BLAST to confirm a unique localization; based on the results, some sequences were further modified and reanalyzed to eliminate potential cross-hybridization to other loci. Potential intra- and intermolecular hybridization of each potential probe sequence were evaluated using the Multiple Primer Analyzer web tool from Thermo Fisher (sensitivity for dimer detection = 3). The resulting candidate probes, consisting of one or two oligonucleotides per repeat, were analyzed experimentally for specific and robust hybridization (Tables 1 and 2).

**Table 1.**
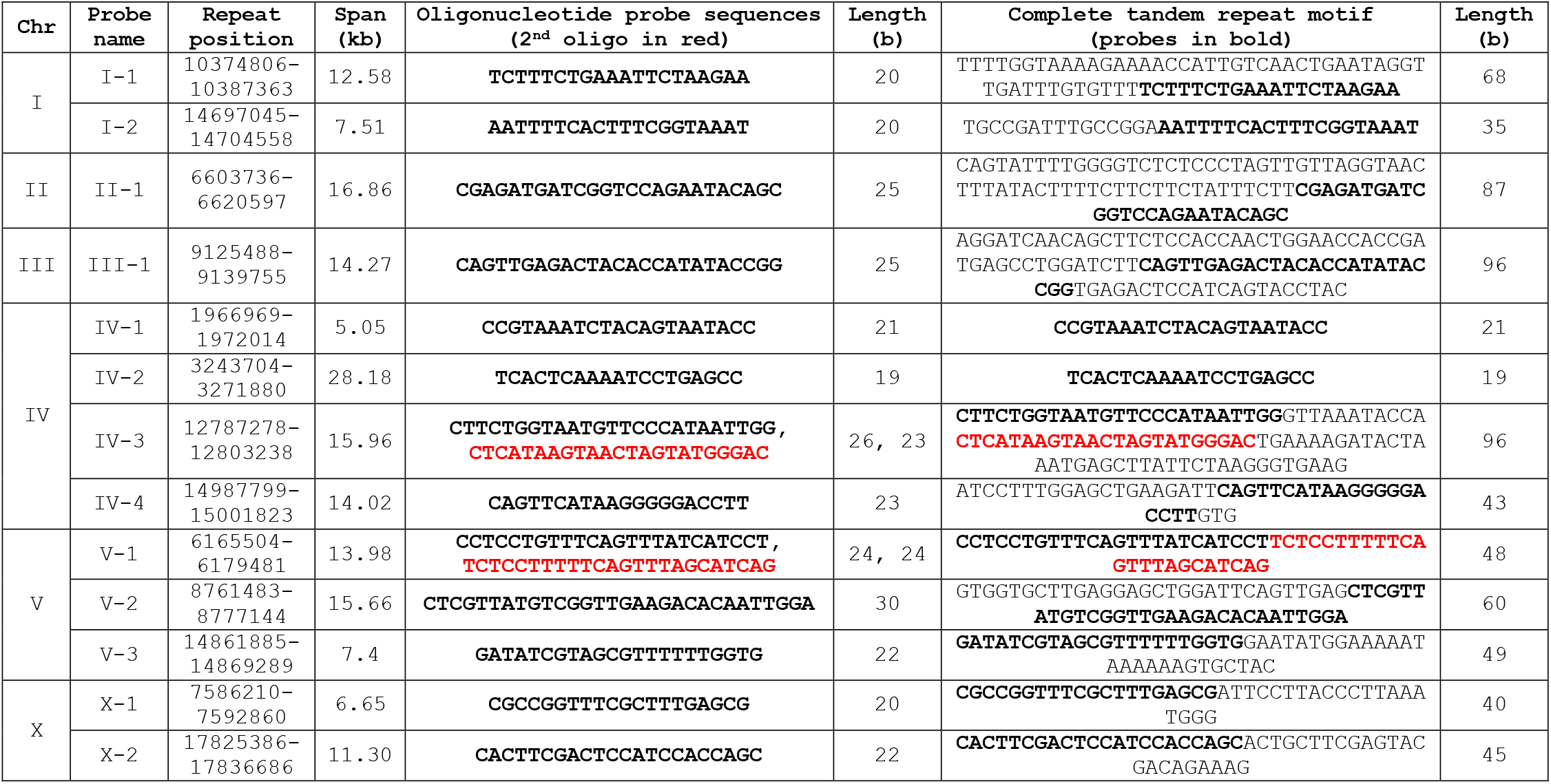
List of locus-specific oligonucleotide FISH probes for *C. elegans*.

**Table 2.**
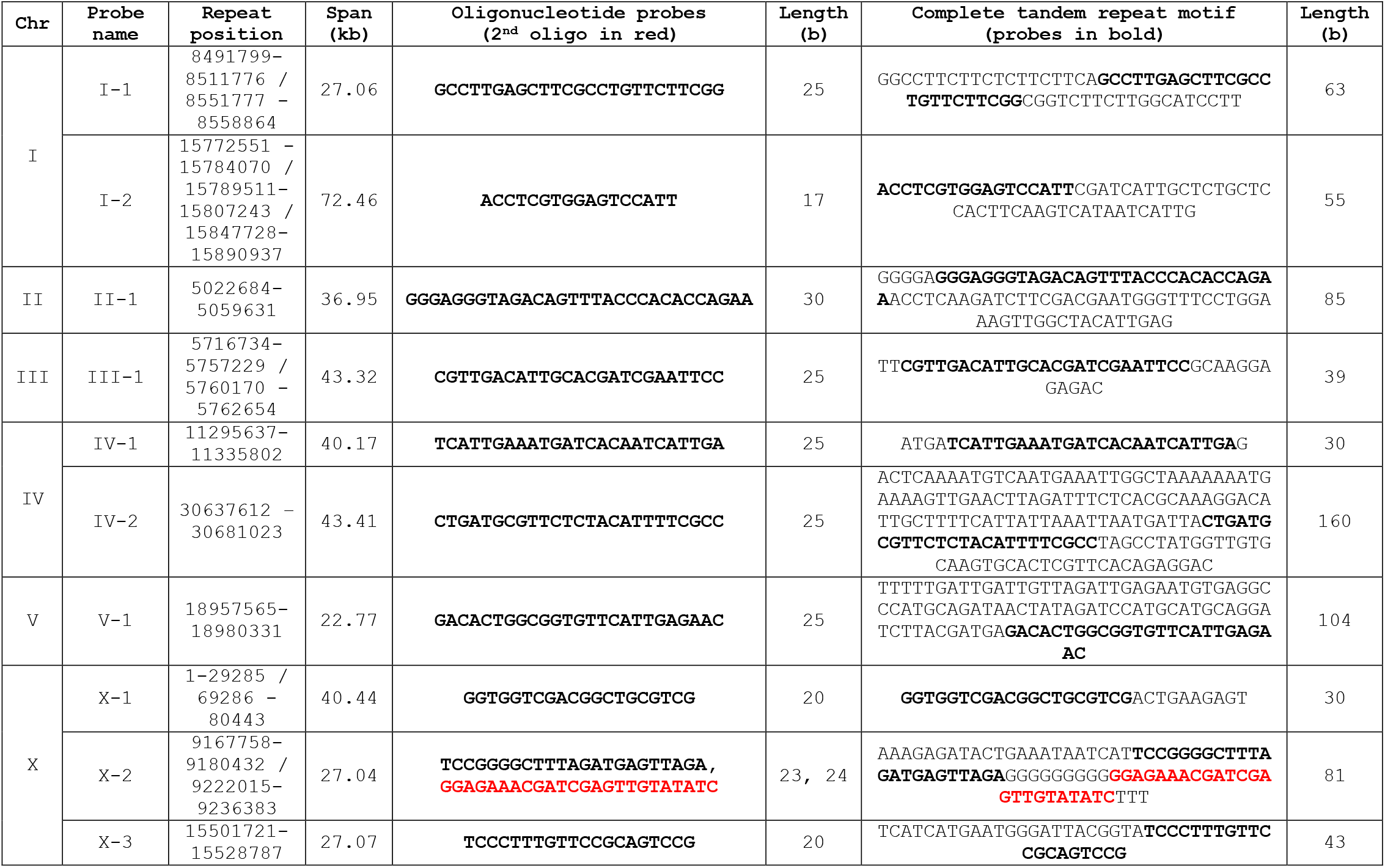
List of locus-specific oligonucleotide FISH probes for *P. pacificus*.

### Oligonucleotide probes

Probes were ordered from IDT as single-stranded DNA oligonucleotides labeled with fluorophores at their 3’ ends (100nM, HPLC purification). To combine different probes in the same experiment, oligos were labeled with Cy3, Cy5, 6-FAM, or Alexa488. Lyophilized oligos were resuspended at 100μM in MilliQ water, aliquoted, and stored at −30°C.

Maps showing the genomic position of each probe (Figure 1A, B) were generated using the R package chromoMap (Anand and Rodriguez Lopez 2022).

**Figure 1.**
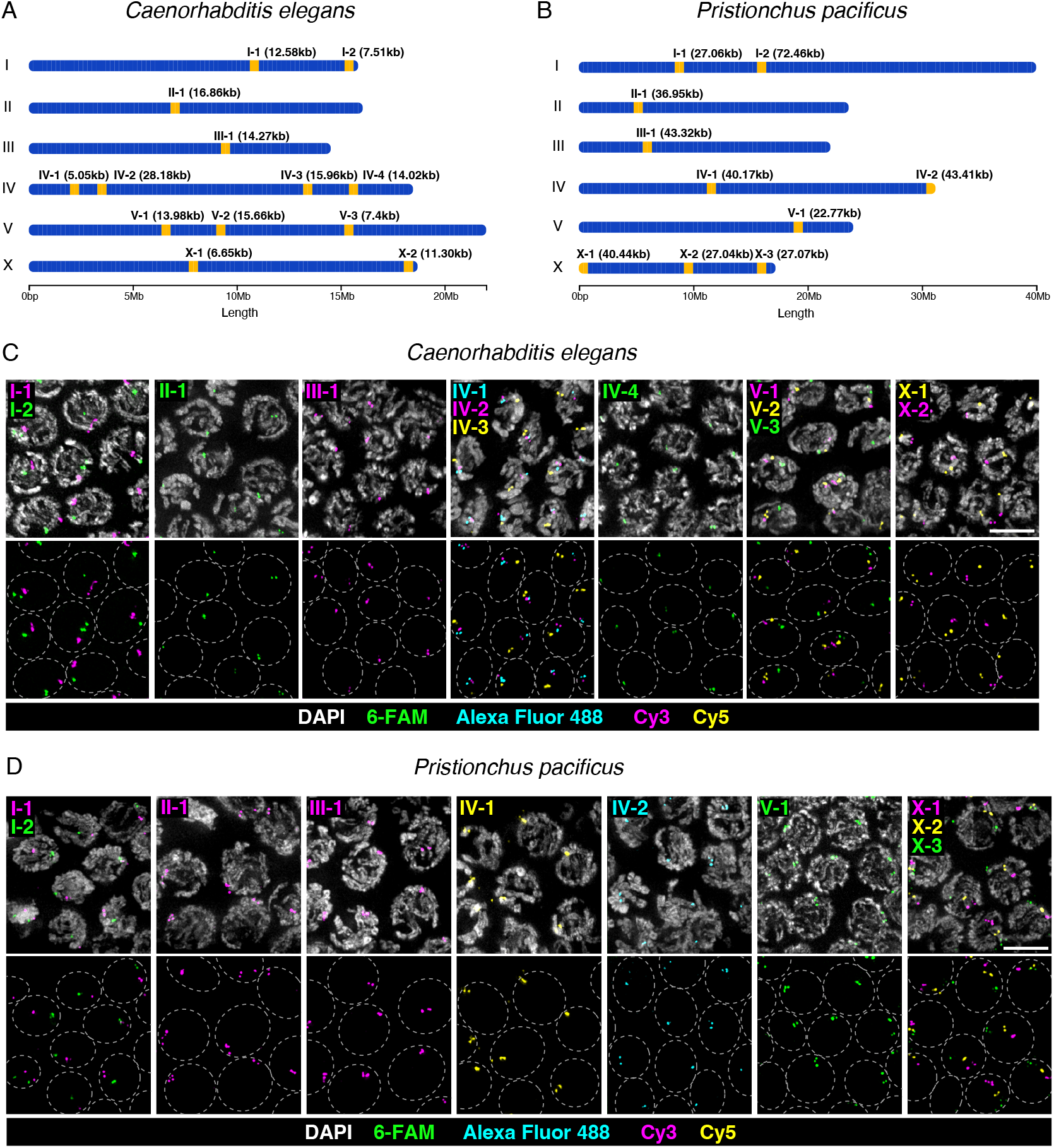
Locus-specific repetitive sequences and FISH probes for *C. elegans* and *P. pacificus*. Chromosome map of oligonucleotide FISH probes for (A) *C. elegans*, and (B) *P. pacificus*; the span of the tandem repeat targeted by each probe is indicated in parentheses. (C, D) Meiotic prophase I nuclei at the pachytene stage after FISH with different combinations of probes. Paired homologous chromosomes display adjacent or overlapping signals for each probe. Gray dashed lines outline the position of the nuclei. All images are maximum-intensity projections of deconvolved 3D stacks. Scale bar = 5μm.

### Fluorescence in situ hybridization and immunofluorescence

Sample preparation and hybridization methods were adapted from prior work on *C. elegans* (Phillips *et al*. 2009) and *P. pacificus* (Rillo-Bohn *et al*. 2021), with minor modifications. Fixed tissue was stained in polypropylene tubes rather than on slides. Briefly, age-matched adult hermaphrodites from either *C. elegans* (N2 and Hawaiian CB4856 strains) or *P. pacificus* (PS312 strain) were transferred to a drop of 30μl of egg buffer (EB) containing 0.05% tetramisole and 0.1% Tween-20 on a coverslip. The gonads were dissected and fixed for 4 minutes by addition of formaldehyde in EB to 2% final, then transferred to a 1.5ml tube containing 1ml PBST. After all samples were dissected and fixed, PBST was replaced with 1ml methanol pre-chilled to −30°C and incubated at room temperature for 5 minutes. Dissected worms were then washed twice with 2x SSCT (0.3 M NaCl, 0.03 M Na citrate, pH 7, 0.1% Tween-20) for 5 minutes and incubated in 200μl 2xSSCT containing 50% formamide solution for 4 hours or overnight at 37°C. Next, the worms were transferred to a 0.2-ml PCR tube, excess solution was removed from the tissue, and 40μl of hybridization mix containing 10-250ng of each probe in hybridization buffer (3x SSC, 48% formamide, 10.6% dextran sulfate) was added. Chromosomal DNA was denatured by incubation in a thermocycler at 91°C for 2 minutes, followed by overnight hybridization at 37°C in the dark. After this incubation, 100μl of 2x SSCT was added to each tube, samples were transferred to a 1.5ml tube, and the tissue was washed three times with 2x SSCT for 5 minutes. After the final wash, the solution was removed and 40μl of SlowFade™ Diamond Antifade Mountant with DAPI (Invitrogen) was added. The tissue was transferred to a slide in a minimal volume of mounting medium, overlaid with a #1.5 high-performance coverslip (Zeiss), and sealed with clear nail polish.

For sequential FISH and immunofluorescence, *C. elegans* and *P. pacificus* expressing endogenously tagged SYP-4::HA proteins (*syp-4(ie29)* (Köhler *et al*. 2020) and *syp-4(ie1002)*, (Rillo-Bohn *et al*. 2021), respectively) were used to visualize the synaptonemal complex. FISH was performed as described above through the second wash after hybridization. The buffer was replaced with 1x Blocking Reagent (Roche) in PBST and the worms were incubated at least 30 min at room temperature. Goat anti-HA (Novus Biologicals #NB600-362) primary antibodies diluted 1:500 into 1X Blocking Reagent (Roche) in PBST were added and incubated overnight at 4°C. Samples were then washed three times with PBST for 5 minutes and incubated with secondary antibodies (1:500 Donkey anti-goat IgG (H+L) conjugated with Cy5 or Cy3, Jackson ImmunoResearch Laboratories, West Grove, PA) for 2 hrs in the dark at room temperature. Finally, the samples were washed three times with PBST for 5 minutes and mounted as described above.

### Microscopy

3D Images were acquired as stacks of optical sections at 0.2μm z-spacing using a DeltaVision Elite microscope (GE) with an Olympus 100X NA 1.45 objective. All data were deconvolved using the constrained iterative algorithm included with the softWoRx package (GE) using 10 cycles and default settings. Maximum-intensity projections were generated from images stacks, and then pseudocolored using Fiji software (Schindelin *et al*. 2012).

### Image analysis

The effect of probe concentration on FISH signal intensity was analyzed using three concentrations of *P. pacificus* Chr X-1 Cy3-labeled probes (0.25ng/μl, 1.25ng/μl, and 6.25ng/μl). The samples were prepared in parallel and imaging conditions and exposure time were the same in each case. Maximum intensity projections of raw images of the Cy3 channel were used for measurements using Fiji (Schindelin *et al*. 2012). The images correspond to the pachytene region of the gonad. A mean background pixel intensity inside the gonad was estimated from the measurement of three ROIs that did not contain FISH signals. This value was subtracted from the images to correct for background. FISH signals were then segmented automatically using the Maximum Entropy thresholding algorithm, which we found empirically to identify fluorescent foci with high accuracy and reproducibility. The mean pixel intensity was measured for each signal and used for subsequent analysis. Statistical analysis and graphing were done using GraphPad Prism (v.9.3.0) software.

## RESULTS

### Identification of tandem repeats and design of locus-specific probes for *C. elegans and P. pacificus*

Our initial analysis of chromosome-level genome assemblies of *C. elegans* and *P. pacificus* using the Tandem Repeat Finder software (Benson 1999) yielded a large number of repeats, 39,505 and 40,651, respectively. After filtering out those with a total span of less than 5kb, we analyzed the remaining candidates using BLAST to confirm a single major cluster in the reference genome (Tables 1 and 2, Figure 1A, B). This yielded 1-4 candidate targets per chromosome for each species.

Thirteen probes for *C. elegans* were characterized. The oligonucleotides used as probes ranged from 19 to 30 bases, and targeted repeats spanning 5.1-28.2 kb (Table 1, Figure 1A). The two tandem repeats on the X chromosome, one near the center and one near the right tip, were previously identified and used successfully as FISH targets, referred as “X center” and “XR” probes, respectively (Phillips *et al*. 2005), while all autosomal targets are described here for the first time. Four additional probes targeting repeats on chromosomes I, II, III, and X showed a smaller unexpected secondary signal during a preliminary FISH test and were excluded from further analysis (Supplementary Table 1). For *P. pacificus*, ten probes were designed and tested; two of these were used to analyze meiotic homolog pairing and synapsis in our recent study (Rillo-Bohn *et al*. 2021). Oligonucleotide probes ranged from 17 to 30 bases, with a target repeat span between 22.8-72.5 kb (Table 2, Figure 1B). None of the repeats we identified are conserved between the two species.

### Probe validation

The performance of the probes for *C. elegans* (N2 strain) and *P. pacificus* (PS312 strain) was primarily analyzed in dissected gonads of adult hermaphrodites, which are densely populated with proliferating germline stem cells (GSCs) and nuclei that span the full range of meiotic stages. At meiotic entry, chromosomes undergo a dramatic transition to form long, individualized territories, and also pair with their homologs. Pachytene nuclei in both species mostly showed two adjacent but discrete signals for each FISH probe, corresponding to the paired homologous loci separated by the synaptonemal complex. Some nuclei displayed up to four smaller signals (Figure 1C, D), likely reflecting separation of the two sister chromatids of each homolog. All of the probes described here produced reliable signals, and none showed unexpected or nonspecific hybridization to secondary loci. Given this very high success rate, additional probes could likely be designed using more relaxed criteria, particularly regarding the total copy number and genomic span of target sequences.

Simultaneous detection of multiple targets in the same sample can be accomplished by combining probes with different fluorescent or hapten labels. We performed double- and triple-labeled hybridizations with sets of probes from the same chromosome (Figure 1C). In *P. pacificus*, three widely-spaced loci on the X chromosome include one from the left end (X-1), one centrally located (X-2), and one near the right end (X-3), their colinear appearance reveals the orientation of the paired chromosomes at the pachytene stage (Figure 1D).

For longer target sequences (i.e., *C. elegans* IV-3 and V-1, and *P. pacificus* X-2), we compared the performance of probes comprising two nonoverlapping oligonucleotides to hybridization with only one of the oligos; surprisingly, the signal intensities with a single probe were quantitatively indistinguishable from data obtained using two labeled oligonucleotides (data not shown).

We also tested two *C. elegans* probes for chromosome I (I-1 and I-2) in the divergent Hawaiian (CB4856) strain. Interestingly, while probe I-1 showed reliable signals in this strain, probe I-2 was not detected (data not shown). This likely reflects strain-specific copy number variation in this repeat; other repeats may also show strain-specific differences and probes should thus be tested before use in strains other than N2/VC2010.

### FISH can be combined with immunolocalization in various tissues

To explore the versatility of the set of probes, we assessed the quality of hybridization in different regions of the gonad and in other tissues of the worms using the same FISH protocol. In the premeiotic region of the *C. elegans* gonad, unpaired loci on homologous chromosomes resulted in two distinct signals, which became closely associated in transition zone nuclei due to chromosome pairing (Figure 2A). Two signals per probe were consistently detected in interphase nuclei of embryos at different developmental stages (Figure 2A).

**Figure 2.**
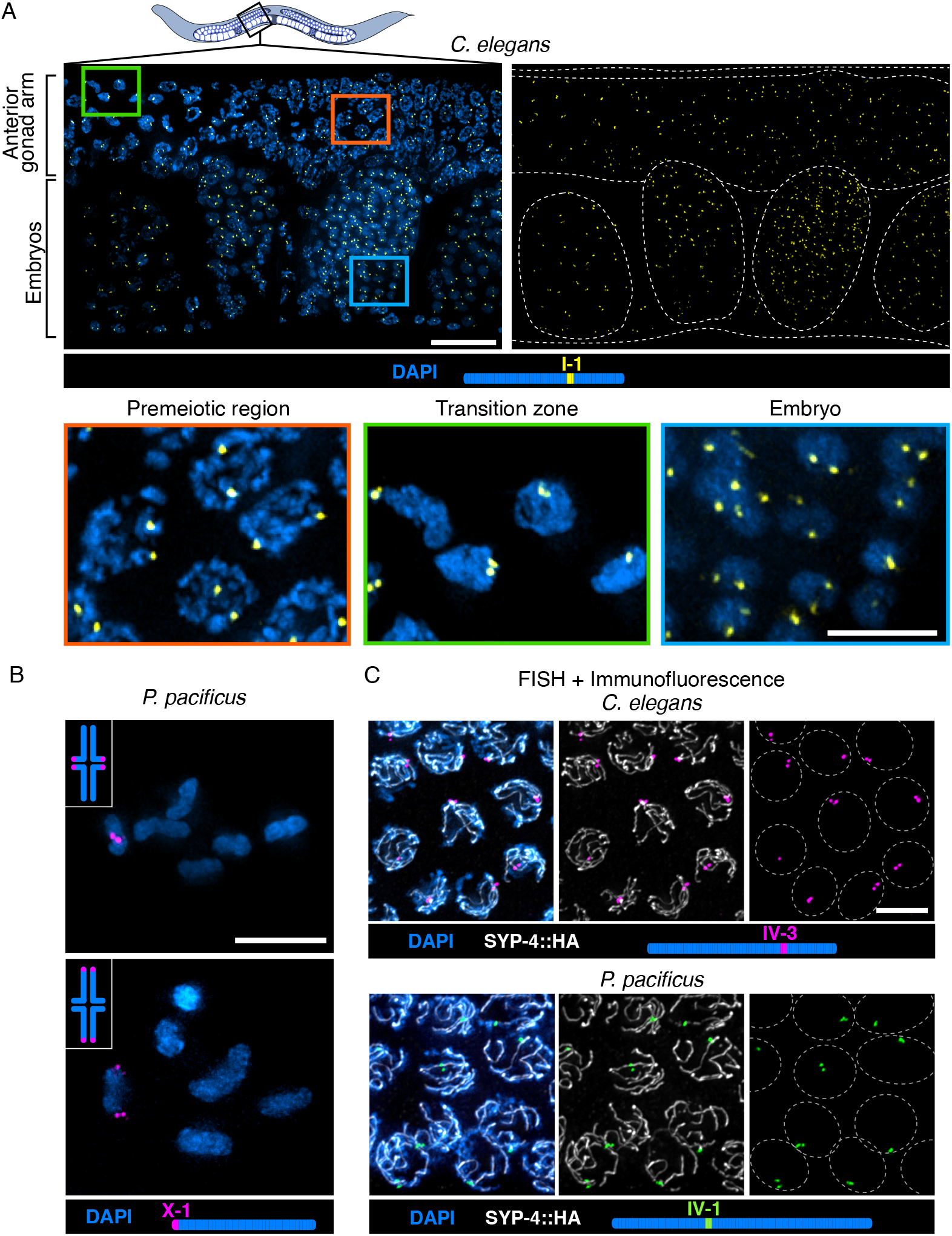
Oligonucleotide FISH probes show robust signals in different tissues and can be combined with immunolocalization. A. Diagram of an adult *C. elegans* hermaphrodite and detail of mid anterior region of the body after FISH with chromosome I-1 probe. The distal anterior gonad arm (top) and four embryos (bottom) show consistent hybridization signals. White dashed lines demarcate the worm body, the gonad arm, and the embryos. Scale bar = 15μm. Higher magnification of nuclei in the premeiotic region with unpaired chromosomes and two distant FISH signals (left), in the transition zone after chromosome pairing (center), and in an embryo (right). Scale bar = 5μm. B. *P. pacificus* diakinesis nuclei showing six bivalents after FISH with chromosome X-1 probe, which reveals chiasma position proximal (top) and distal (bottom) to the left of the X chromosome end. Scale bar = 5μm. C. FISH followed by immunolocalization of the synaptonemal complex protein SYP-4 in strains of *C. elegans* and *P. pacificus* expressing SYP-4::HA. Scale bar = 5μm. Worm diagram adapted from Wikimedia Commons *Caenorhabditis elegans hermaphrodite adult-en.svg* by K. D. Schroeder, CC-BY-SA 3.0. All images are maximum-intensity projections of deconvolved 3D stacks.

In *C. elegans* and *P. pacificus*, the position of the single crossover between each pair of homologous chromosomes stochastically determines their orientation during the first meiotic division; the end farther from the crossover site usually separates from its homolog earlier but retains sister chromatid cohesion and leads toward the spindle pole during the first division (Albertson and Thomson 1993; Rillo-Bohn *et al*. 2021). This specification of a “long arm” and a “short arm” along each bivalent is mediated by asymmetric disassembly of axis and synaptonemal complex proteins during diplotene-diakinesis (Martinez-Perez *et al*. 2008; Rillo-Bohn *et al*. 2021). This can be demonstrated using FISH probes that detect distal chromosome regions. For example, hybridization to *P. pacificus* gonads with the X-1 probe reveals that this locus can be on either the long arm or the short arm at diakinesis (Figure 2B; see also Albertson and Thomson 1993, Rillo-Bohn *et al*. 2021).

In situ hybridization to repetitive targets is advantageous for experiments involving localization of both DNA and proteins because it is relatively tolerant of different fixation methods, allowing optimization of immunofluorescence. We sequentially combined the FISH protocol with immunolocalization of the synaptonemal complex protein SYP-4 in strains of both species expressing HA-tagged SYP-4. This combined protocol resulted in well-preserved morphology and high signal/background for both targets (Figure 2C). For antigens that are highly sensitive to overfixation, immunolocalization can be performed after an initial light fixation, followed by post-fixation to further stabilize the tissue before hybridization (Phillips *et al*. 2009).

### Oligonucleotide probes show consistent hybridization signals over a wide range of concentrations

To determine the amount of probe required for detection of a typical repetitive locus, we tested three different concentrations of *P. pacificus* X-1 probe over a 25-fold dilution range (i.e., 0.25ng/μl, 1.25ng/μl, and 6.25ng/μl). All of these conditions resulted in bright and consistent hybridization signals with low background in the gonad (Figure 3A). After measuring the background-corrected mean pixel intensity of each FISH signal at the pachytene stage, the mean FISH signal intensity using the intermediate concentration (1.25ng/μl) was significantly higher than the other two concentrations. This likely reflects minor variations in conditions other than probe concentration, since no significant difference was detected between the extremes of the range (0.25ng/μl and 6.25ng/μl) (Figure 3B). Moreover, we observed similar gonad-to-gonad variation within the same sample as between samples (Figure 3C). This analysis is consistent with our prior experience that hybridization to repetitive targets is fairly insensitive to variations in probe concentration, and that low probe concentrations are sufficient to detect repetitive targets spanning 5 kb or more with robust signal/background ratios.

**Figure 3.**
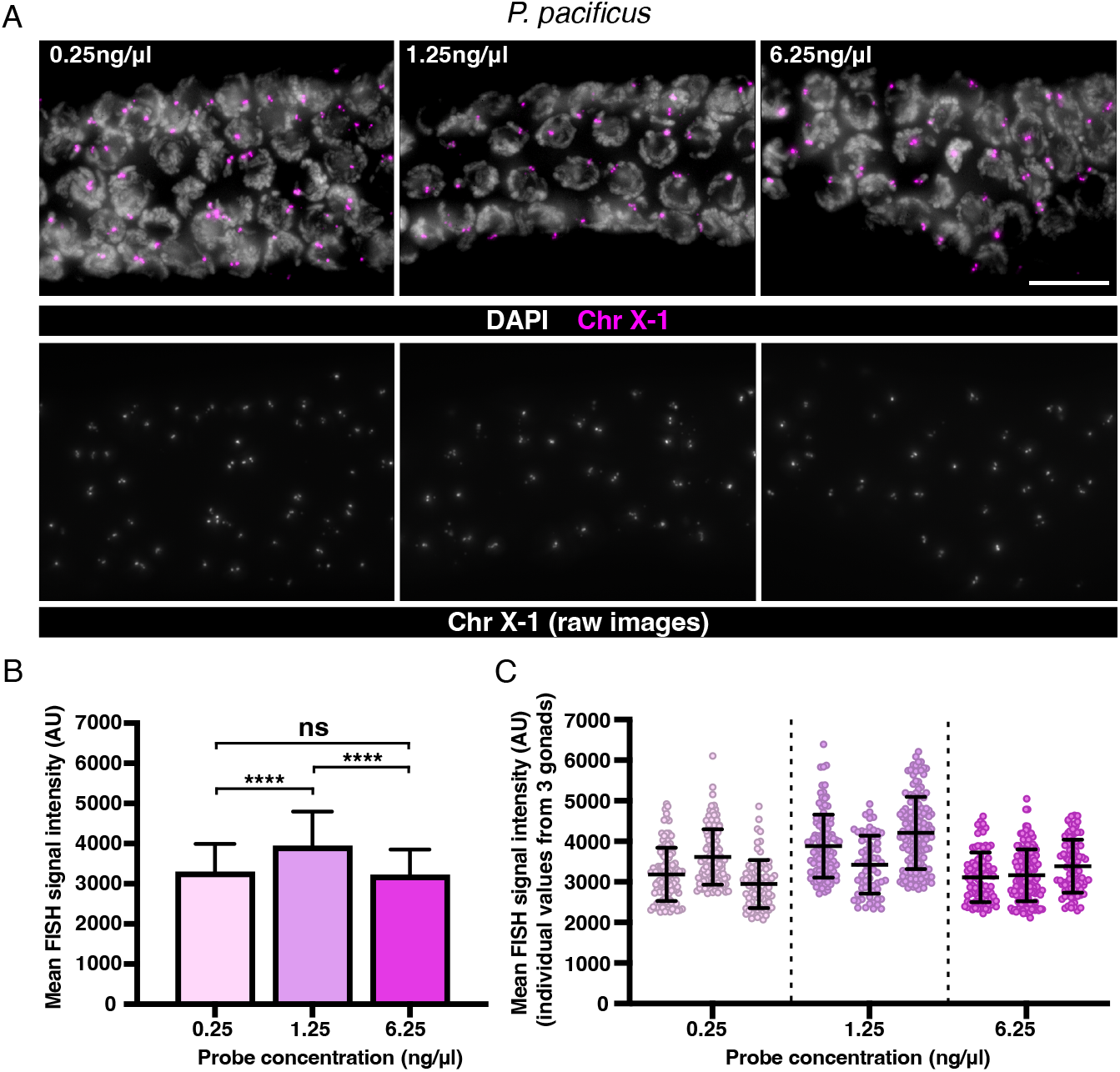
Comparison of signal intensities resulting from hybridization with different probe concentrations. A. Maximum-intensity projections of raw (not deconvolved) 3D data showing the pachytene region of *P. pacificus* gonads after FISH with Cy3-labeled X-1 probes at 0.25ng/μl (10ng), 1.25ng/μl (50ng), and 6.25ng/μl (250ng). Representative images of Cy3 channel used for image analysis. Scale bar = 10μm. B. Graph showing the background-corrected FISH signal intensity for each concentration of X-1 probe (mean + SD, ****p<0.0001 by Welch’s t-test). C. Disaggregated data from the three images measured per concentration, showing the inter-specimen variability (mean ± SD).

## DISCUSSION

We describe a simple method to design locus-specific DNA FISH probes targeting tandem repeats in model and non-model organisms. The main requirement for this approach is a chromosome-level genome assembly that includes accurate annotation and quantification of repetitive elements. Using this procedure, we validated probes for the nematodes *C. elegans* and *P. pacificus*, including a least one marker for each chromosome. These probes show reliable hybridization, and their synthesis as short oligonucleotides (17-30 bases) facilitates diffusion within different fixed tissues, including intact embryos. This protocol can also be combined with immunolocalization or other cytological methods.

To maximize convenience and reproducibility, probes can be commercially synthesized as labeled oligonucleotides. Given that the lowest concentration we tested (0.25ng/μl, or 10ng in 40μl of hybridization mix) resulted in signal/background indistinguishable from a 25-fold higher concentration, a small-scale commercial synthesis of a fluorescently labeled probe with HPLC purification, yielding ~2 nMol of product, provides sufficient material for at least 2,000 hybridizations. The methods presented here can be easily adapted to generate cost-effective probes in any species with a complete genome assembly based in part on long-read sequencing.

## ACKNOWLEDGMENTS

We thank members of the Dernburg lab for helpful discussions during the course of this work.

---

**Supplementary Table 1.**
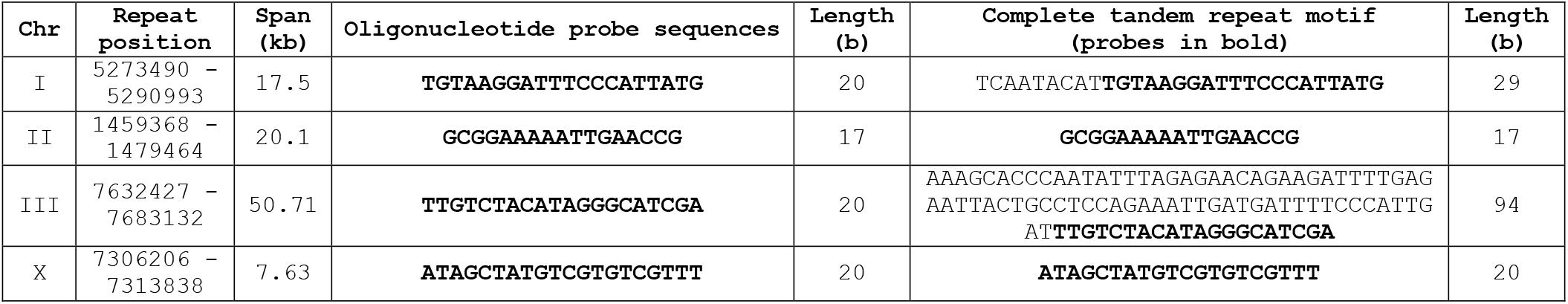
List of oligonucleotide FISH probes for *C. elegans* with extra signals.

## LITERATURE CITED

Albertson D. G., 1984 Localization of the ribosomal genes in *Caenorhabditis elegans* chromosomes by in situ hybridization using biotin-labeled probes. The EMBO Journal 3: 1227–1234. https://doi.org/10.1002/j.1460-2075.1984.tb01957.x

Albertson D. G., and J. N. Thomson, 1993 Segregation of holocentric chromosomes at meiosis in the nematode *Caenorhabditis elegans*. Chromosome Res 1: 15–26. https://doi.org/10.1007/BF00710603

Anand L., and C. M. Rodriguez Lopez, 2022 ChromoMap: an R package for interactive visualization of multi-omics data and annotation of chromosomes. BMC Bioinformatics 23: 33. https://doi.org/10.1186/s12859-021-04556-z

Bauman J. G. J., J. Wiegant, P. Borst, and P. van Duijn, 1980 A new method for fluorescence microscopical localization of specific DNA sequences by in situ hybridization of fluorochrome-labelled RNA. Experimental Cell Research 128: 485–490. https://doi.org/10.1016/0014-4827(80)90087-7

Beliveau B. J., E. F. Joyce, N. Apostolopoulos, F. Yilmaz, C. Y. Fonseka, et al., 2012 Versatile design and synthesis platform for visualizing genomes with Oligopaint FISH probes. Proceedings of the National Academy of Sciences 109: 21301–21306. https://doi.org/10.1073/pnas.1213818110

Benson G., 1999 Tandem repeats finder: a program to analyze DNA sequences. Nucleic Acids Research 27: 573–580. https://doi.org/10.1093/nar/27.2.573

Dernburg A. F., J. W. Sedat, and R. S. Hawley, 1996 Direct evidence of a role for heterochromatin in meiotic chromosome segregation. Cell 86: 135–146.

Dernburg A. F., K. McDonald, G. Moulder, R. Barstead, M. Dresser, et al., 1998 Meiotic recombination in *C. elegans* initiates by a conserved mechanism and is dispensable for homologous chromosome synapsis. Cell 94: 387–398.

Ferreira H. C., B. D. Towbin, T. Jegou, and S. M. Gasser, 2013 The shelterin protein POT-1 anchors *Caenorhabditis elegans* telomeres through SUN-1 at the nuclear periphery. Journal of Cell Biology 203: 727–735. https://doi.org/10.1083/jcb.201307181

Heckmann S., J. Macas, K. Kumke, J. Fuchs, V. Schubert, et al., 2013 The holocentric species *Luzula elegans* shows interplay between centromere and large-scale genome organization. Plant J 73: 555–565. https://doi.org/10.1111/tpj.12054

Köhler S., M. Wojcik, K. Xu, and A. F. Dernburg, 2020 The interaction of crossover formation and the dynamic architecture of the synaptonemal complex during meiosis. bioRxiv. https://doi.org/10.1101/2020.02.16.947804.

Martinez-Perez E., M. Schvarzstein, C. Barroso, J. Lightfoot, A. F. Dernburg, et al., 2008 Crossovers trigger a remodeling of meiotic chromosome axis composition that is linked to two-step loss of sister chromatid cohesion. Genes & Development 22: 2886–2901. https://doi.org/10.1101/gad.1694108

Nguyen P., K. Sahara, A. Yoshido, and F. Marec, 2010 Evolutionary dynamics of rDNA clusters on chromosomes of moths and butterflies (Lepidoptera). Genetica 138: 343–354. https://doi.org/10.1007/s10709-009-9424-5

Pardue M. L., and J. G. Gall, 1969 Molecular hybridization of radioactive DNA to the DNA of cytological preparations. Proceedings of the National Academy of Sciences 64: 600–604. https://doi.org/10.1073/pnas.64.2.600

Phillips C. M., K. L. McDonald, and A. F. Dernburg, 2009 Cytological Analysis of Meiosis in *Caenorhabditis elegans*, pp. 171–195 in Meiosis, Methods in Molecular Biology. edited by Keeney S. Humana Press, Totowa, NJ.

Rillo-Bohn R., R. Adilardi, T. Mitros, B. Avşaroğlu, L. Stevens, et al., 2021 Analysis of meiosis in *Pristionchus pacificus* reveals plasticity in homolog pairing and synapsis in the nematode lineage. eLife 10: e70990. https://doi.org/10.7554/eLife.70990

Rödelsperger C., J. M. Meyer, N. Prabh, C. Lanz, F. Bemm, et al., 2017 Single-Molecule Sequencing Reveals the Chromosome-Scale Genomic Architecture of the Nematode Model Organism *Pristionchus pacificus*. Cell Reports 21: 834–844. https://doi.org/10.1016/j.celrep.2017.09.077

Rödelsperger, Christian, Streit, Adrian, and Sommer, Ralf J, 2013 Structure, Function and Evolution of The Nematode Genome, in eLS, John Wiley & Sons, Ltd: Chichester.

Ruiz-Ruano F. J., M. D. López-León, J. Cabrero, and J. P. M. Camacho, 2016 High-throughput analysis of the satellitome illuminates satellite DNA evolution. Sci Rep 6: 28333. https://doi.org/10.1038/srep28333

Schindelin J., I. Arganda-Carreras, E. Frise, V. Kaynig, M. Longair, et al., 2012 Fiji: an open-source platform for biological-image analysis. Nat Methods 9: 676–682. https://doi.org/10.1038/nmeth.2019

Schmidt T. L., B. J. Beliveau, Y. O. Uca, M. Theilmann, F. Da Cruz, et al., 2015 Scalable amplification of strand subsets from chip-synthesized oligonucleotide libraries. Nat Commun 6: 8634. https://doi.org/10.1038/ncomms9634

Yoshimura J., K. Ichikawa, M. J. Shoura, K. L. Artiles, I. Gabdank, et al., 2019 Recompleting the *Caenorhabditis elegans* genome. Genome Res. 29: 1009–1022. https://doi.org/10.1101/gr.244830.118

